# Over-eruption in marsupial carnivore teeth: compensation for a constraint

**DOI:** 10.1101/2023.03.16.533044

**Authors:** Menna E. Jones

## Abstract

Pronounced over-eruption of the canine teeth, much greater than in eco-morphologically equivalent placental carnivores, occurs with age and growth in Australian marsupial carnivores. Suppression of functional tooth replacement is a characteristic of marsupials and is frequent among diverse placentals, where the primitive therian pattern is two generations of incisor, canine and premolar teeth. Rapid determinate growth is one among multiple hypotheses proposed to explain the loss of tooth replacement in mammals. In this line of reasoning, the animal reaches a sufficient body or jaw size to accommodate adult-sized teeth by the time it requires functional dentition. Marsupial carnivores have a full set of adult anterior teeth at weaning, which erupt into a juvenile jaw that is 25 to 30 % adult size, compared with well grown in placental carnivores. Indeterminate over-eruption of the canine teeth in marsupial carnivores results in increasing canine height and diameter with increasing body size with age, suggesting this is a compensatory mechanism for the constraint of a single generation of anterior teeth. Patterns of over-eruption in different tooth types of marsupial carnivores are consistent with two non-exclusive mechanisms that operate in other mammals, a response to tooth wear and lack of an occlusal partner.

## Introduction

There has been long interest in comparing the biology of placental and marsupial mammals, and how phylogenetically distinct traits relate to adaptations and convergent eco-morphological niches in different parts of the world (e.g. 1, 2). One trait of interest is the ontogenetic pattern of tooth replacement, with marsupials and some placental mammals exhibiting the monophyodont condition and most placentals the diphyodont condition. The mammalian pattern of two generations of anterior dentition is considered to relate to lactation and rapid determinant growth (3). In this primitive therian pattern of diphyodont tooth replacement, no teeth are required during early lactation, one set of small deciduous teeth scaled to the size of the juvenile jaw are fully erupted by weaning, and these are replaced by a successional and permanent larger set of incisors, canines and premolars as the animal approaches its mature body size. Molar teeth are not replaced as these erupt sequentially at the rear of the tooth row as the animal grows (3, 4). Rapid determinate growth in mammals mitigates the need for the continual tooth replacement seen in non-mammal vertebrates with indeterminate growth (3). Vestigial first generation anterior teeth, the incisors and canines, are characteristic of marsupial mammals (5) including the carnivorous marsupials in the Family Dasyuridae (6, 7), and variable suppression is frequent among diverse taxa of placental mammals (3). In species with this monophyodont condition, the logic follows that the jaw will be sufficiently large to accommodate adult-sized teeth at weaning when the animal needs functional dentition (3).

This logic of adult-sized teeth scaled to jaw size at weaning is not supported in the case of the Australian marsupial carnivores (Order Dasyuromorphia, Families Thylacinidae and Dasyuridae) which have monophyodont tooth replacement. In these taxa, the single set of anterior teeth, which must serve the animal as a fully grown adult, erupt fully (enamel cervical margin level with gum line and no root exposed) into a jaw that is much too small to accommodate them (4). The single set of incisors and canines are the second generation successors to the rudimentary non-erupting first generation teeth, and the two deciduous anterior premolars (P1 and P2) are relatively developmentally retarded with no evidence of successional replacements (5, 7). Only in the third premolar position (P3), in the primitive condition, does both a deciduous and successional tooth erupt (5, 6, 8, 9), although even P3 replacement appears to be lacking in some marsupial carnivores (10). One set of molars erupt as in placentals (8). The question then is how the pattern of tooth replacement in marsupial carnivores affects ontogenetic dental function.

The pattern of tooth replacement in marsupial carnivores may operate as a phylogenetic constraint if the fully erupted anterior teeth of marsupial carnivores are smaller than the anterior successional teeth of eco-morphologically equivalent placental carnivores at the same developmental stage. Comparative analyses that would reveal whether this is the case are precluded, however, because measures of canine size are affected by canine shape, which influences tooth function and varies with killing and hunting type (2), and so are confounded with phylogeny. Werdelin (4) explored the consequences of the marsupial pattern of tooth replacement for the sequence of tooth eruption, jaw growth and molar morphology. He suggested that the pattern of tooth replacement does impose a constraint on tooth and jaw morphology during ontogeny, and could result in major differences in evolutionary plasticity. Post-birth marsupial teat attachment has been proposed as an explanation for mechanical suppression of odontogenesis in marsupials (4). The widespread suppression of tooth replacement in placental mammals, however, begs alternative explanations (3).

Pronounced over-eruption, greater than in eco-morphologically similar placental carnivores, is evident in the canine teeth of all extant and recently extinct marsupial carnivore species. Tooth over-eruption in mammals is reported to be a response to both tooth wear and to lack of an occlusal partner to that tooth (11, 12). The effects of these mechanisms are not mutually exclusive. If tooth wear is a factor, teeth with occlusal partners (molar teeth) should remain the same height as tooth wear progresses. Teeth without complete occlusion (canine teeth) should increase in height and over-erupt to a greater extent, both in response to tooth wear and the lack of an occlusal partner.

In this paper I explore whether over-eruption in canine teeth in Australian marsupial carnivores functions as a compensation for the constraint placed on dental ontogeny by the suppression of anterior tooth replacement. Specifically, does over-eruption of the unreplaced canine teeth result in an increase in tooth length or diameter that may compensate for an initially smaller tooth erupting into a small juvenile jaw? I consider the Australian carnivorous marsupial taxa in the Order Dasyuromorphia: the thylacine (*Thylacinus cynocephalus*) in the Family Thylacinidae, and the Tasmanian devil (*Sarcophilus harrisii*) and four species of quolls (*Dasyurus maculatus, D. viverrinus, D. geoffroyi* and *D. hallucatus*) in the Family Dasyuridae. The smaller, largely insectivorous, members of the Dasyuridae are not considered.

## Methods

Tooth measurements were taken from wild animals with repeated measures through time as the animals aged and tooth and skull measurements from museum specimens. Museum specimens of Australian marsupial carnivores were measured in Australian museum collections: the Tasmanian Museum and Art Gallery (Hobart, Tasmania), the Queen Victoria Museum (Launceston, Tasmania), the Museum of Victoria (Melbourne, Victoria), the Donald Thomson and the Department of Fisheries and Wildlife Collections housed at the Museum of Victoria, the Australian Museum (Sydney, NSW), the Macleay Museum of the University of Sydney, the Queensland Museum (Brisbane, Queensland) and the South Australian Museum in Adelaide. Museum specimens of placental carnivores were measured in the collection of the Tel Aviv Museum of Natural History at Tel Aviv University in Israel. Measurements of live, wild-living devils, spotted-tailed quolls and eastern quolls were collected from a healthy population free of devil facial tumour disease (DFTD) during field work in the northern part of the Cradle Mountain - Lake St. Clair National Park in Tasmania, Australia. All measurements were taken with vernier callipers (0.01 mm accuracy).

### Measurements

Tooth measurements were defined as follows: 1) Canines. Total height was measured on skulls from the midline of the crest of the alveolar bone to the tip on skulls, and in live animals from the midline of the crest of the gum or from a clear brown stain indicating that position where there was obvious gum recession; crown height from the midline or crest of the enamel cervical margin (the edge of the enamel that covers the crown of the tooth) to the tip (Figure 1); maximum anteroposterior (APD) and mediolateral (MLD) diameters corresponded to the diameters at the level of the alveolar crestal bone (skulls) or gum line (live animals). 2) Molars. Total height was measured from the crest of the alveolar bone to the tip of the tallest cusp; crown height from the crest of the enamel cervical margin to the tip of the tallest cusp by holding the calipers level with the margin and crest on both anterior and posterior roots simultaneously (Figure 1). Extent of over-eruption for all teeth, defined as the height of exposed root above the dentine-enamel margin (ERH), was calculated for both canines and molars as the difference between tooth and crown height. I follow the nomenclature for the molar teeth (M1-M4) commonly used by Australian zoologists (13).

**Figure 1.**
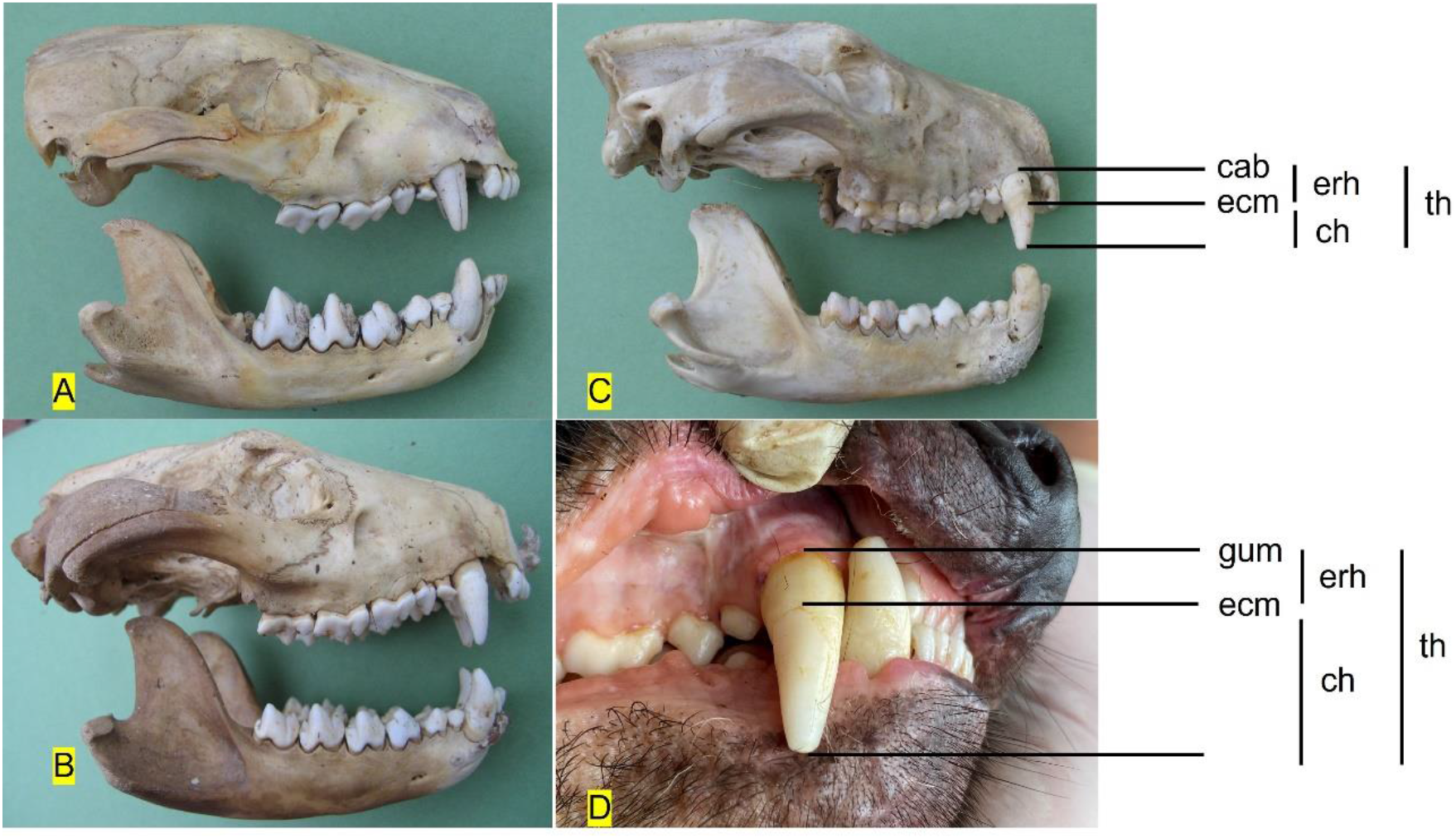
Images of the teeth of Tasmanian devils showing canine tooth measurements, and over-eruption and wear in canine and molar teeth at different ages. A) Juvenile at weaning age - first three molars fully erupted and a cusp appearing on the lower but not yet on the upper M4; tooth wear minimal (score 1) on the canines and M1, and no wear on M2 and M3. B) Juvenile in subadult or first year of independence (1-2 years old with four molars fully erupted and no over-eruption evident in canines or molars; tooth wear minimal (score 1) on the canines and M1, and no wear on M2–M4. C) Older devil (approximately 4 year old) with over-eruption pronounced on canine (enamel cervical margin half-way down the tooth) and evident in lower first molar M1; tooth wear substantial with scores of 3 with wear facets on upper and lower canines, and 4, 3, 3, 2 for M1-M4. D) Two year old wild male devil with some over-eruption and wear (score 2) on the canine. A-C from museum specimens; D wild devil (Photo: XX). cab = crest of alveolar bone, gum = gum line, ecm = enamel cervical margin, erh = exposed root height, ch = crown height, th = total height. Tooth wear scores: 0 = no wear, tip or cusp sharp; 1 = tip or cusp worn, no dentine exposed; 2 = tip or cusp worn, dentine exposed; 3 = tip or cusp well worn, dentine exposed, less than half tooth height worn; 4 = tip or cusp well worn, dentine exposed, more than half tooth height worn; 5 = tooth worn to gum.

One upper canine tooth and the lower molar set on one side of the jaw (M1-M4) were measured on each marsupial skull or wild animal and one upper canine tooth on the skulls of placentals. Wear stage of all teeth measured was recorded using schemes devised for canines of marsupial carnivores (14, 15) and for lower molars of devils (Pemberton, 1990): 0 = no wear, tip or cusp sharp; 1 = tip or cusp worn, no dentine exposed; 2 = tip or cusp worn, dentine exposed; 3 = tip or cusp well worn, dentine exposed, less than half tooth height worn; 4 = tip or cusp well worn, dentine exposed, more than half tooth height worn; 5 = tooth worn to gum. The canine tooth wear criteria were suitable and recorded also for placental carnivores.

Body size was measured as condylobasal length (CBL, from the base of I1 to the tip of the occiput) and jaw length (JAW, from the base of I1 to the condyle on the mandible) on all skulls. Jaw length was used in analyses of museum material where the high incidence of broken skulls markedly reduced sample sizes, as mandibles have lower incidents of breakage. Body size in wild animals was represented by measurement of pes length but was not used in analyses or figures. Pes length, measured from the heel to the end of the metacarpals with the phalanges bent over, is a precise measure of size in live marsupial carnivores (14).

### Museum collections

In museum collections, all substantially undamaged skulls were measured, with teeth broken in life (some wear on the broken edge) or following death (sharp edges to the break) excluded from measurement. Only wild-caught individuals were sampled; captive-sourced individuals were excluded from the study. Samples of measured teeth from all species included a range of tooth wear stages, indicating that the samples were not biased towards young or old individuals. Individuals that were small juveniles were excluded from analysis.

Wild-caught specimens were measured for all six extant and recently extinct Australian species of marsupial carnivores (Table 1, Figure 2): the thylacine, devil, spotted-tailed quoll, eastern quoll, western quoll and northern quoll, although sample sizes sufficient for statistical analyses were obtained for the first four species only. Total height (from tip to the alveolar crest), crown height, and the two maximum diameters of canine teeth (antero-posterior and medio-lateral), and total height and crown height of molar teeth were measured. All four teeth in the molar row were measured for devils and thylacines, and just M1 and M4 for spotted-tailed and eastern quolls. Teeth of individuals ranging in age and size from independent juveniles (defined from field observations as M4 commenced but not completed eruption) to fully grown adults (defined as complete dental rupture and a combination of size and tooth wear, as sutures do not close in adults) were measured.

**Table 1.**
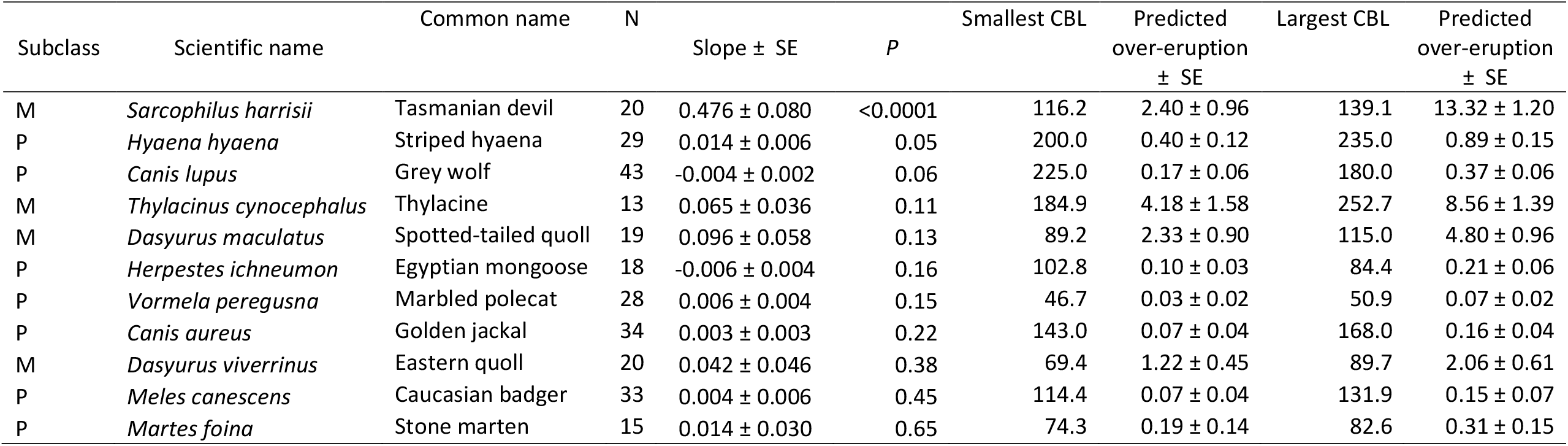
The relationship between the amount of over-eruption and skull length (CBL) in marsupial and placental carnivores with predicted values of over-eruption for the smallest and largest individual of each species in the data set. The “estimate” is the slope of the regression for each species. CBL and extent of over-eruption are measured in mm. Subclasses: M = marsupial; P = placental. N = number of individuals. CBL = condylobasal length measured on the skull. SE = standard error.

| Subclass | Scientific name | Common name | N | Slope ± SE | *P* | Smallest CBL | Predicted over-eruption ± SE | Largest CBL | Predicted over-eruption ± SE |
| --- | --- | --- | --- | --- | --- | --- | --- | --- | --- |
| M | *Sarcophilus harrisii* | Tasmanian devil | 20 | 0.476 ± 0.080 | <0.0001 | 116.2 | 2.40 ± 0.96 | 139.1 | 13.32 ± 1.20 |
| P | *Hyaena hyaena* | Striped hyaena | 29 | 0.014 ± 0.006 | 0.05 | 200.0 | 0.40 ± 0.12 | 235.0 | 0.89 ± 0.15 |
| P | *Canis lupus* | Grey wolf | 43 | -0.004 ± 0.002 | 0.06 | 225.0 | 0.17 ± 0.06 | 180.0 | 0.37 ± 0.06 |
| M | *Thylacinus cynocephalus* | Thylacine | 13 | 0.065 ± 0.036 | 0.11 | 184.9 | 4.18 ± 1.58 | 252.7 | 8.56 ± 1.39 |
| M | *Dasyurus maculatus* | Spotted-tailed quoll | 19 | 0.096 ± 0.058 | 0.13 | 89.2 | 2.33 ± 0.90 | 115.0 | 4.80 ± 0.96 |
| P | *Herpestes ichneumon* | Egyptian mongoose | 18 | -0.006 ± 0.004 | 0.16 | 102.8 | 0.10 ± 0.03 | 84.4 | 0.21 ± 0.06 |
| P | *Vormela peregusna* | Marbled polecat | 28 | 0.006 ± 0.004 | 0.15 | 46.7 | 0.03 ± 0.02 | 50.9 | 0.07 ± 0.02 |
| P | *Canis aureus* | Golden jackal | 34 | 0.003 ± 0.003 | 0.22 | 143.0 | 0.07 ± 0.04 | 168.0 | 0.16 ± 0.04 |
| M | *Dasyurus viverrinus* | Eastern quoll | 20 | 0.042 ± 0.046 | 0.38 | 69.4 | 1.22 ± 0.45 | 89.7 | 2.06 ± 0.61 |
| P | *Meles canescens* | Caucasian badger | 33 | 0.004 ± 0.006 | 0.45 | 114.4 | 0.07 ± 0.04 | 131.9 | 0.15 ± 0.07 |
| P | *Martes foina* | Stone marten | 15 | 0.014 ± 0.030 | 0.65 | 74.3 | 0.19 ± 0.14 | 82.6 | 0.31 ± 0.15 |

**Figure 2.**
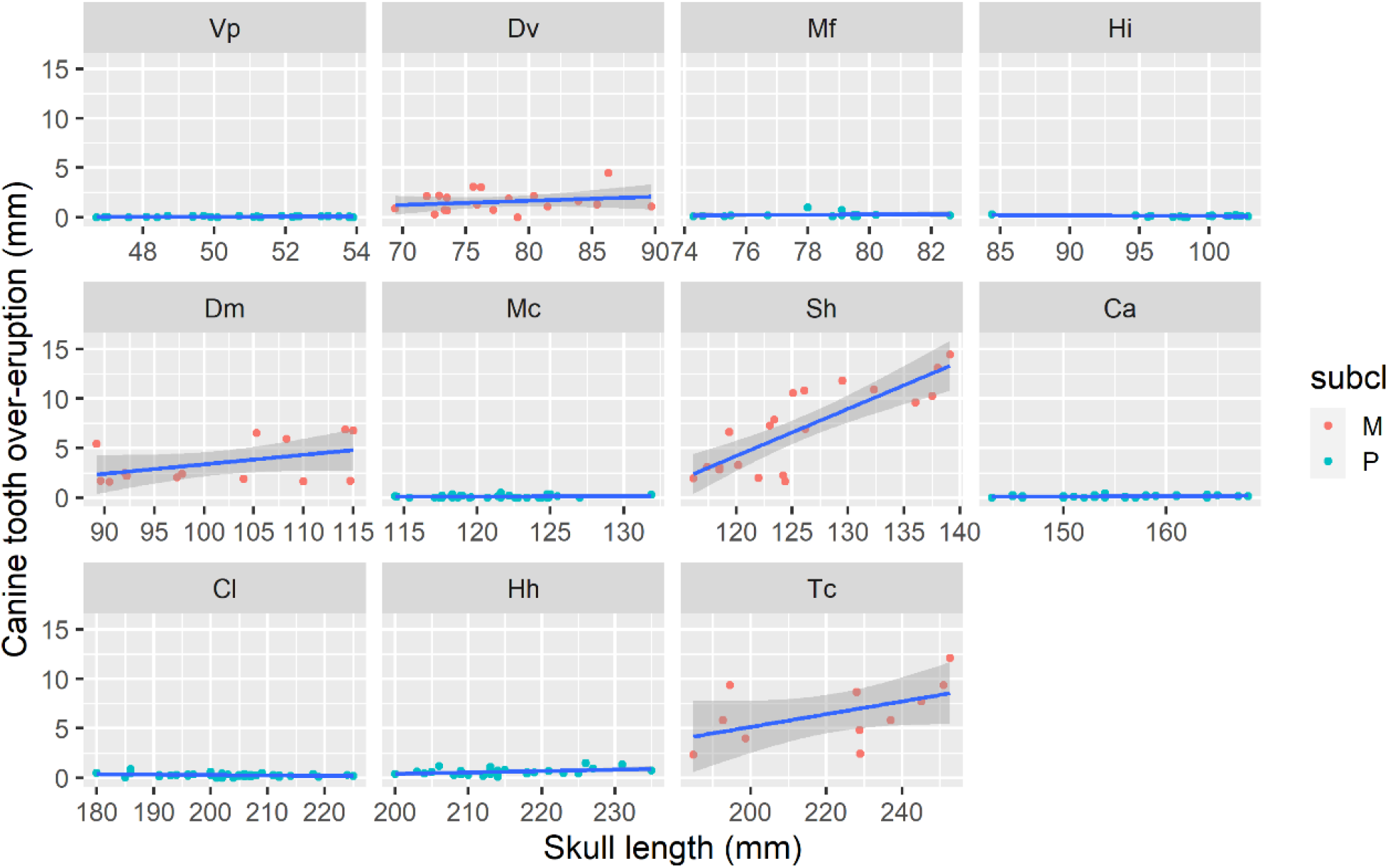
The extent of over-eruption of the canine teeth plotted against skull length (CBL) for adult animals four species of marsupial carnivores and six species of similar-sized and ecologically similar placental carnivores held in museum collections. Species are presented left to right, top to bottom, from smallest to largest skull size. All measurements are in mm. subcl = mammalian subclass, with “M” red symbols = marsupial species and “P” blue symbols = placental species. Symbols are first letter of genus and species name. Marsupial carnivores: Dv = *D. viverrinus*, eastern quoll; Dm = *Dasyurus maculatus*, spotted-tailed quoll; Sh = *Sarcophilus harrisii*, Tasmanian devil; Tc = *Thylacinus cynocephalus*, thylacine. Placental carnivores: Vp = Vormela peregusna, marbled polecat; Mf = *Martes foina*, beech marten; Hi = *Herpestes ichneumon*, Egyptian mongoose; Mc = *Meles canescens*, Caucasian badger; Ca = *Canis aureus*, golden jackal; Cl = *Canis lupus*, grey wolf; Hh = *Hyaena hyaena*, spotted hyaena.

Total height and crown height of canine teeth were measured on skulls of wild-caught adult specimens (complete dental rupture and closure of sutures) of seven species of placental carnivores (Table 1, Figure 2). All species are native to Israel and thus were well represented in the museum collection. These species represent the diversity of placental families that are eco-morphologically equivalent to the Australian marsupial carnivore species, with species of similar body size selected for study (2). The wolf (*Canis lupus*) and the jackal (*Canis aureus*) are convergent with and larger and smaller, respectively, than the thylacine. The striped hyaena (*Hyaena hyaena*) is larger than but eco-morphologically similar to the devil, while the badger (*Meles meles*) is the same size as the devil. The Egyptian mongoose (*Herpestes ichneumon*) and the stone marten (*Martes foina*) are eco-morphological analogues of the marsupial quolls (2).

### Wild animals

For devils, spotted-tailed quolls and eastern quolls, the total height, crown height and the two maximum diameters of the canine teeth and pes length were measured repeatedly during growth on wild individuals trapped over a two-year period. The age of all individuals in the data set were known definitively because these individuals had been initially trapped in the population as independent juveniles. Time since birth and independence were recorded. Note that time since birth in marsupial carnivores includes time spent in the pouch following a 3-week gestation, and time in a den. Time since independence was standardised as the number of months since February (for devils) or the previous December (for spotted-tailed quolls) in the year of independence. Timing of independence, assessed by regression of mammae in the adult female population, is highly synchronised in all three species and occurs in late January in healthy populations of devils free of DFTD (16), in December in spotted-tailed quolls 0 (17) and by November in the eastern quoll (18).

### Statistical analyses

The relationship between the amount of canine over-eruption and body size across marsupial and placental carnivore species in museum collections was analysed using generalised linear models (GLM), with a gaussian distribution and species and log(CBL) as covariates. Changes in canine tooth dimensions over time in wild marsupial carnivores were analysed using a generalised linear mixed model using a likelihood ratio test (LRT, chi-squared) to test for significance. Species comparisons among marsupial carnivores in the changes in three canine dimensions in museum specimens were analysed using a GLM with gaussian distribution and species and jaw length as covariates. To compare the trends of over-eruption or height across the four-tooth molar row (from M1 to M4) with body size amongst species, the many-model function in the Broom package was used (https://r4ds.had.co.nz/many-models.html; accessed 23 December 2022). These models were fit with linear, quadratic and cubic trend functions.

Throughout this paper, *n* is the number of observations, and *r*^*2*^ *adj* is the adjusted squared multiple *r*. All analyses were conducted in the R Project for Statistical Computing (version 4.2.2, 2022.10.31) using R Studio (version 2022.12.0).

## Results

Skulls and canine teeth for the comparison of canine over-eruption between marsupial and placental carnivores were measured from between 13 and 43 individuals per species in museum collections (Table 1). For analysis of canine tooth eruption and dimensions in marsupial carnivores, the number of different individuals measured on museum specimens and wild animals were: thylacine (23 museum, 0 wild), Tasmanian devil (33 museum, 153 wild individuals with repeat measurements through time of canine teeth), spotted-tailed quoll (20 museum, 19 wild), eastern quoll (20 museum, 61 wild). Sample sizes for analysis of over-eruption and height of molar teeth, only from museum specimens, ranged from 40 in each species of quoll to 88 in thylacines and 124 in devils (Table 3). The much higher natural and post-mortem fracture rates of sabre-like canine teeth compared to more rounded molar teeth account for the discrepancy in sample sizes from museum specimens between canine and molar teeth. Variation in sample sizes amongst different measures reflect tooth breakage or loss in individual wild or museum specimens. Data were also obtained for western quoll (4 museum, 0 wild), and northern quoll (3 museum, 0 wild) but sample sizes were insufficient for statistical analysis.

### Over-eruption of canine teeth

There are clearly different slopes amongst species of marsupial and placental carnivores when over-eruption is regressed on body size (skull size; CBL) (interaction between body size and CBL: F10, 244 = 18.27, p < 0.0001; museum specimens). All marsupial carnivore species show more pronounced over-eruption than any placental carnivore species across the whole range of body sizes measured (Table 1, Figure 2). Over-eruption significantly increased with body size for Tasmanian devils and marginally so for striped hyaenas but not in other species (Table 1). These two species are the only two specialised bone-eaters or osteophages in the data set.

### Canine tooth dimensions increase with body size and age

In marsupial carnivores measured in museum collections, canine tooth size increased with body size (jaw length) in three dimensions (total length, antero-posterior and medio-lateral diameters) for all four species (eastern quolls, spotted-tailed quolls, Tasmanian devils, thylacines) (Figure 3). The linear trends of these relationships varied among species for canine length and antero-posterior diameter with less evidence for differences among species for medio-lateral diameter (Table 2). Eastern quolls (intercept) had the steepest increase in canine size with log(jaw length) on all three measures. For canine length, the slopes were a bit shallower for Tasmanian devils, more so for spotted-tailed quolls and much shallower for thylacines. For antero-posterior diameter, the slopes were a bit shallower for Tasmanian devils, more so for thylacines and much shallower for spotted-tailed quolls. For medio-lateral diameter, spotted-tailed quolls trended towards a shallower slope, although overall the slopes were not different (Table 2, Figure 3).

**Table 2.** Parameter estimates of the differences among species (species by body size interaction) in the slope of the relationship between canine tooth dimensions (length, antero-posterior and medio-lateral diameters) and body size (jaw length) for museum specimens of four species of marsupial carnivores. EQ = eastern quoll, SQ = spotted-tailed quoll, TD = Tasmanian devil and TH = thylacine. SE = standard error, t – t test, *p* = p value.

| Measure | Species | Slope | SE | t | $p$ |
| --- | --- | --- | --- | --- | --- |
| Canine length |  |  |  |  |  |
|  | EQ (intercept) | 1.745 | 0.241 | 7.239 | <0.001*** |
|  | SQ | -0.758 | 0.274 | -2.767 | 0.007** |
|  | TD | -0.635 | 0.277 | -2.295 | 0.024* |
|  | TH | -1.079 | 0.274 | -3.945 | <0.001*** |
| Antero-posterior diameter |  |  |  |  |  |
|  | EQ (intercept) | 1.616 | 0.180 | 8.991 | <0.001*** |
|  | SQ | -0.823 | 0.205 | -4.028 | <0.001*** |
|  | TD | -0.428 | 0.206 | -2.075 | 0.041* |
|  | TH | -0.538 | 0.204 | -2.638 | 0.01** |
| Medio-lateral diameter |  |  |  |  |  |
|  | EQ (intercept) | 1.298 | 0.196 | 6.624 | <0.001*** |
|  | SQ | -0.464 | 0.223 | -2.082 | 0.041* |
|  | TD | -0.206 | 0.225 | -0.917 | 0.362 |
|  | TH | -0.213 | 0.222 | -0.956 | 0.342 |

**Figure 3.**
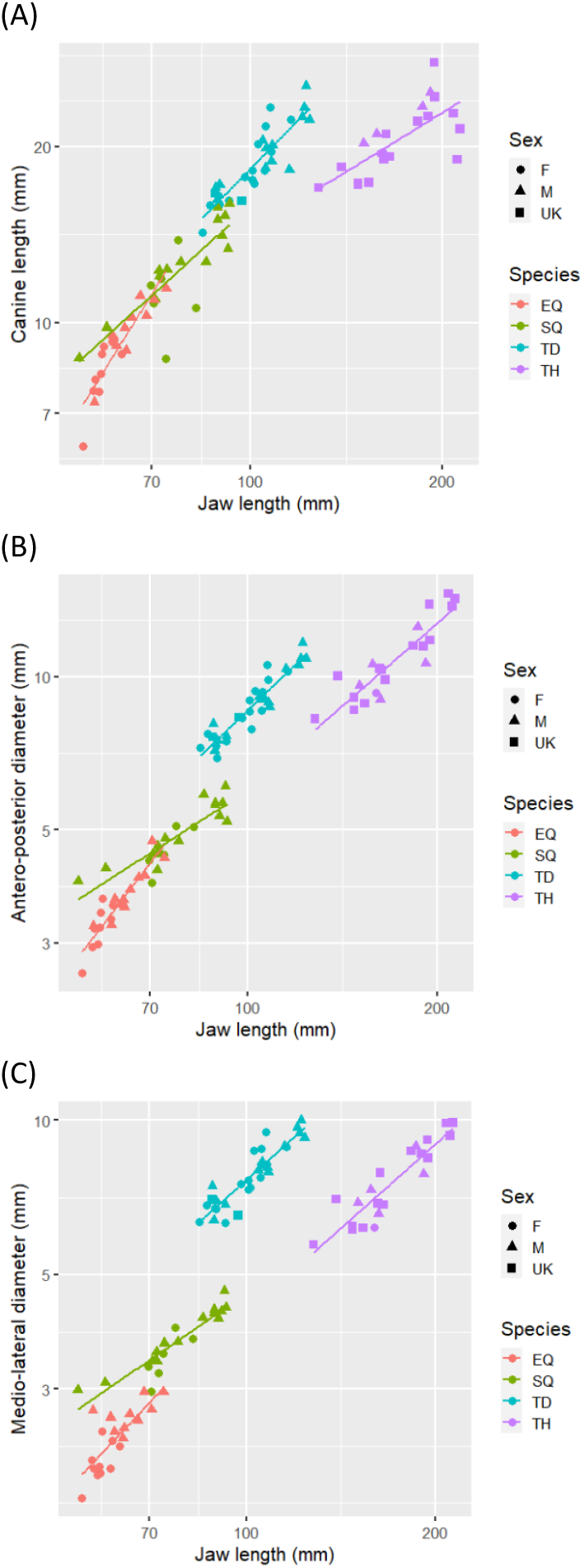
Changes in canine dimensions with increasing body size (jaw length) in four wild-captured Australian dasyuromorph carnivores from museum collections: total canine length (A); antero-posterior diameter (B); and medio-lateral diameter (C). All measurements in mm. canine length = right canine length, measured from the apex of the alveolar crest to the tip; the two canine tooth diameters measured at the skull edge. JAW = jaw or mandible length. The individual figure titles include species code and individual identification number. EQ = eastern quoll, SQ = spotted-tailed quoll, TD = Tasmanian devil and TH = thylacine.

In wild marsupial carnivores canine tooth size increased through time in three dimensions (total length, antero-posterior and medio-lateral diameters) (CBL: LRT = 12.67, p <0.0001; APD: LRT = 16.09, p <0.0001; MLD: LRT = 22.78, p <0.0001) but with no effect of the animal’s sex (CBL: LRT = 0.24, p = 0.62; APD: LRT = 0.56, p = 0.46; MLD: LRT = -0.42, p = 1.00) (Figure S1, Supplementary). There was variation amongst species and individuals, with pronounced increases in the single spotted-tailed quoll and most Tasmanian devils and little or no changes on all dimensions in eastern quolls, and on some dimensions in Tasmanian devils. For the spotted-tailed quoll and the Tasmanian devils, the individuals whose canine teeth increased in length and the two diameters were independent juveniles at the first capture and were growing in body size over the time period of the measurements. The total age since birth and months since weaning for these individuals were: the spotted-tailed quoll (SQ7 - 7 months/4 months) and Tasmanian devils: TD117 (11 months/2 months), TD120 (13 months/4 months), TD 54 (11 months/2 months), TD 59 (11 months/2 months), TD65 (11 months/2 months), TD69 (11 months/2 months) and TD75 (13 months/4 months). Note that time since birth in marsupial carnivores includes time spent in the pouch following a 3-week gestation, and time in a den. Other Tasmanian devils were already mature or old at the commencement of the study and had completed growth, examples being TD15 and TD45 (both age 3 years at first measurement). In these two old individuals, while canine length did not change or decreased slightly over time, the two canine diameters continued to increase. While the canine teeth continue to over-erupt with age and the conical tooth becomes wider, tooth wear is an ongoing process eroding the tip of the tooth and reducing the total length of the tooth in old animals (Figure 1).

### Over-eruption of molar teeth

The extent of over-eruption in each molar tooth (M1 to M4) by body size (jaw length) was described by a linear trend in museum specimens of all four species of marsupial carnivores for which there were sufficient sample sizes (Figure 4a). The interaction term testing for differences in these slope coefficients across the molar tooth row (M1 to M4) indicated that these within-tooth linear trends became less steep across the tooth row. In all four species, over-eruption was more pronounced in M1 and least in M4. This pattern was highly significant for devils, significant for thylacines, approaching significance for eastern quolls, and not significant for spotted-tailed quolls (Table 3a). This means that for devils and thylacines, as the individual grows the amount of over-eruption is highest in M1 and sequentially less for M2, M3 and M4 and this happens in a linear fashion (Figure 4a). This shows that in these larger species, over-eruption in the molar teeth scales with body size according to their eruption sequence. The teeth that erupt prior to weaning when the animal is small show a lot of over-eruption between the time of their eruption and when the individual reaches adult size. This phenomenon becomes progressively less along the molar tooth row so that M4, which erupts five months after weaning in Tasmanian devils, shows little over-eruption.

**Table 3.** Coefficients for the linear trends of the relationship between a) molar over-eruption and b) molar height and jaw length for each tooth, M1 to M4, across the molar tooth row for marsupial carnivores in museum collections. EQ = eastern quoll, SQ = spotted-tailed quoll, TD = Tasmanian devil and TH = thylacine. SE = standard error, t – t test, *p* = p value, Radj = adjusted r^2^, N = sample size.

| Species | Linear trend | SE | t | $p$ | Radj | N |
| --- | --- | --- | --- | --- | --- | --- |
| a) Molar over-eruption and jaw length |  |  |  |  |  |  |
| TD | -0.02982 | 0.009307 | -3.20343 | 0.001754 | 0.724293 | 124 |
| SQ | -0.00545 | 0.009697 | -0.56169 | 0.577805 | 0.598754 | 40 |
| EQ | -0.02363 | 0.013506 | -1.74994 | 0.088649 | 0.569291 | 40 |
| TH | -0.01076 | 0.004421 | -2.43459 | 0.017135 | 0.821568 | 88 |
| b) Molar height and jaw length |  |  |  |  |  |  |
| EQ | -0.04126 | 0.018342 | -2.24947 | 0.030692 | 0.183943 | 40 |
| SQ | -0.02828 | 0.015788 | -1.79096 | 0.082206 | 0.280759 | 38 |
| TD | -0.00637 | 0.015476 | -0.41187 | 0.68127 | 0.477064 | 114 |
| TH | 0.02278 | 0.007481 | 3.045123 | 0.003195 | 0.830572 | 84 |

**Figure 4.**
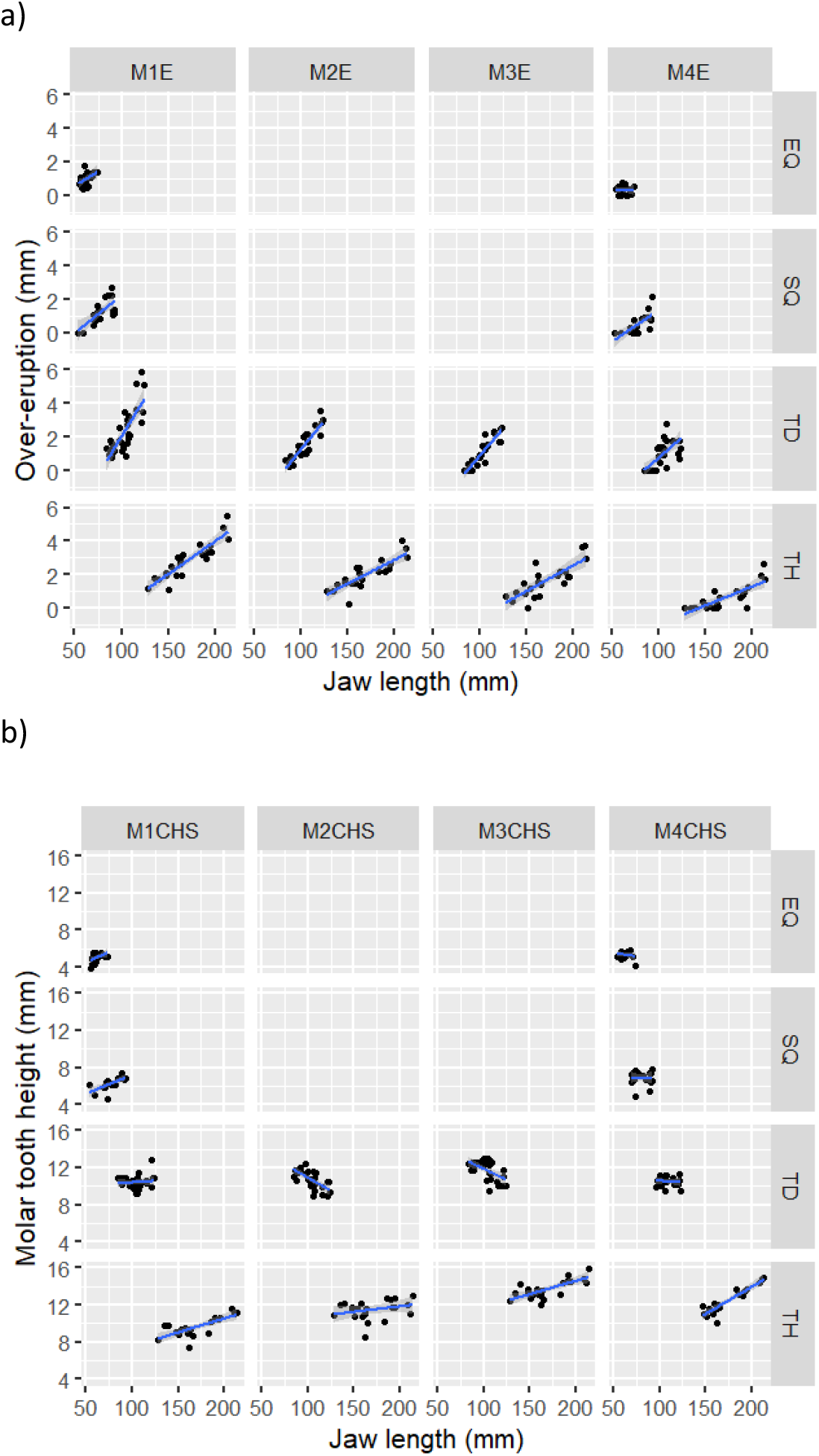
Extent of a) over-eruption and b) tooth height across the molar tooth row regressed against jaw length in marsupial carnivores from museum collections. EQ = eastern quoll, SQ = spotted-tailed quoll, TD = Tasmanian devil and TH = thylacine.

The trend in molar tooth height with increasing body size (jaw length) in museum specimens was best described by a quadratic function for thylacines and Tasmanian devils. The trend was, by definition, linear for eastern and spotted-tailed quolls because only two teeth were measured (M1 and M4). In thylacines, molar tooth height increased significantly with body size (jaw length; which reflects increasing age) for all four molar teeth (Figure 4b). The interaction between linear trends for each tooth was significant and positive, with later erupting molar teeth taller and increasing more in height than earlier erupting teeth (Table 3b, Figure 4b). The greatest increase in height with body size (jaw length) was in M4, then in M1 and M3, and least in M2 (Figure 4b). In Tasmanian devils, the interaction amongst these slope coefficients along the molar tooth row (M1 to M4) was not significant (Table 3b), indicating that molar height does not increase as the animal gets larger and older; indeed M2 and M3 decrease in height (Figure 4b). The interaction between the linear trend in molar height with increasing body size between M1 and M4 was significant for eastern quolls and approaching significance for spotted-tailed quolls (Table 3b). In both species, molar height increased with size and age in M1 but not in M4 (Table 3b, Figure 4b), according to the eruption sequence.

### Composite patterns of over-eruption and tooth height in relation to tooth wear

Figure 5 shows the composite patterns of total tooth height, including the contribution of over-eruption, with increasing tooth wear for canine and molar teeth in repeatedly measured individuals of Tasmanian devils, spotted-tailed quolls and eastern quolls in a wild population. For all three species, these stacked bar graphs show a general pattern of increasing canine length but stable or slightly decreasing molar height with wear stage. In both canine and molar teeth, there is an increasing contribution to total tooth height from over-eruption with increasing tooth wear. When the same measures were plotted for museum specimens, these patterns were not evident (Figure S2, Supplementary). The reason for this difference is probably that each data point in the museum data set represented a different individual, compared with repeated measures over time of the same individuals in the wild data set. Tooth wear and over-eruption are expected to increase with time and age. The wild data set will reflect this temporal component. The museum data set reflects snapshots in time. Canine teeth, in particular, can remain sharp (low wear score), though, even with a significant amount of over-eruption, as evident in devils and male spotted-tailed quolls (Figure S2).

**Figure 5.**
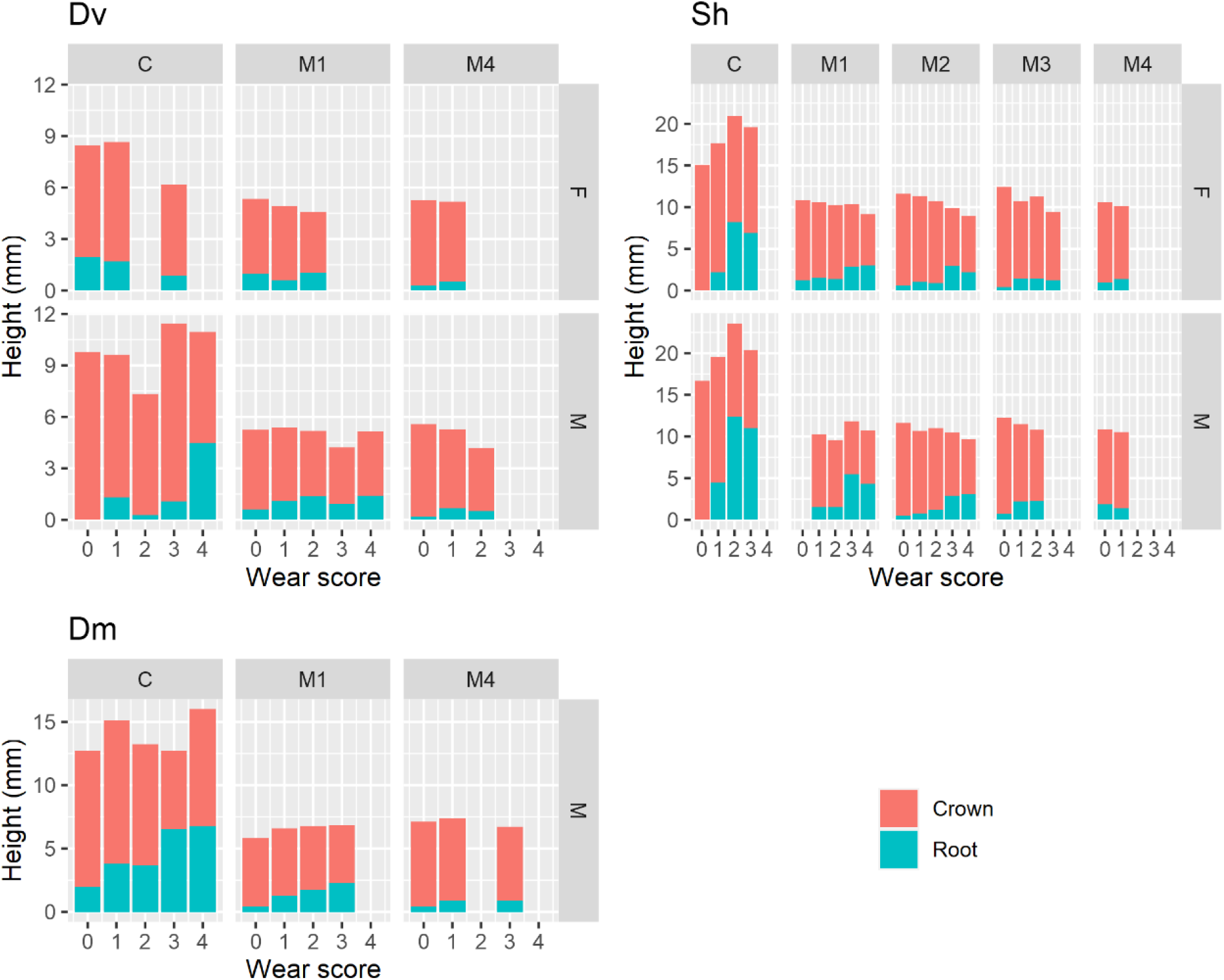
Canine and molar tooth height for each tooth wear stage in marsupial Tasmanian devils, spotted-tailed quolls and eastern quolls, wild-caught at Cradle Mountain National Park in Tasmania, Australia. The stacked bars separate the components of total tooth height that comprise the enamelled crown and the exposed root where the tooth has over-erupted. Tooth types are: canine (C), and molars 1 through 4 (M1, M2, M3, M4). Wear stages are 0 (no wear), 1 (tip or cusp worn), 2 (tip or cusp worn, dentine exposed), 3 (tip or cusp well worn, dentine exposed, less than half tooth height worn), 4(tip or cusp well worn, dentine exposed, more than half tooth height worn). F = female. M = male. There were too few data to analyse female spotted-tailed quolls.

## Discussion

Over-eruption of the canine teeth is evident in placental carnivores, but it is so marked in marsupial carnivores that in some, presumably very old individuals with advanced tooth wear, the enamel cervical margin has progressed to near the tip of the canine. This suggests that canine teeth in these marsupial carnivores erupt continuously through life. While dynamic erosion of the alveolar bone at the crest of where the canine tooth emerges from the maxilla may explain some of the variation in the increasing height of the canine tooth with age, the differences between marsupial and placental carnivores and the extreme over-eruption in old animals suggests this is not a general explanation. This phenomenon of canine over-eruption is not restricted to the dasyuromorph marsupial carnivores. Continuous growth of canine teeth well into adult life or throughout life has been reported in extinct South American marsupial carnivore lineages: the “dog-like” borhyaenids (Family Borhyaenoidea) (19, 20), and in the marsupial saber-tooth carnivores (Family Thylacosmilidae) (21). In these species, the canine teeth remained open-rooted, which is not the case in dasyuromorph marsupials.

Is pronounced over-eruption of the canines a compensatory mechanism for the potential constraint imposed by eruption of the single set of anterior dentition into a juvenile-sized jaw that must also serve the individual as an adult? Juvenile devils and quolls possess a fully erupted set of anterior teeth at independence (weaning), when they are only a quarter to a third of their final adult size (approximate body weights: eastern quoll - 0.35kg at weaning, adult female 1kg, adult male 1.5kg; spotted-tailed quoll - 1kg at weaning, female 2.5kg, male 4kg; and devil at weaning 2.5kg, female 7.5kg, male 10kg, respectively) (unpublished data and 17). In contrast, placental carnivores are well grown when their second generation or successional anterior teeth replace the small first set present at weaning (4). Canine tooth size is a three-dimensional combination of crown to tip length and antero-posterior and medio-lateral diameters, which relates tooth function to killing behaviour and hunting type and is confounded with phylogeny (2, 22-24). This complicates direct comparisons of tooth size relative to body size between marsupial and placental carnivores. If over-eruption is a compensatory mechanism, however, the single generation of canine teeth of marsupial carnivores at weaning should be small and scaled to the size of the jaw (and body size); the marsupial canine teeth should increase in size, probably in both height and diameter, as the individual grows; and the phenomenon of over-eruption should occur to a greater extent than in similar-sized placental carnivores.

Over-eruption of canine teeth in marsupial carnivores results in a net increase in height and diameter as the individual grows to adult size, with the greatest rate of increase coinciding with the period of rapid growth from small juvenile at independence to adult size. Inspection of canine teeth removed from the mandible or jaw, particularly those of devils which have robust canines that are almost circular in cross-section that are adaptations for bone-eating (2, 24), reveal the maximal width is about half-way along the tooth which tapers conically (Figure 1) at both ends. Presumably even in very old animals, the distal half of the tooth which diminishes in diameter remains within the skull or jaw. The conical shape of the proximal half of the canine tooth ensures a gradual increase in both the length and width of the canine teeth with over-eruption. This over-eruption and increasing canine tooth size occurs during growth, so that the weaned juvenile has small canine teeth scaled to its’ smaller skull and jaw size and the fully grown adult has large canine teeth scaled to its’ size. The rate of increase in canine dimensions is generally scaled with body size of the species, being greatest in the smallest species, the eastern quoll, and least in the thylacine. The devil is an exception to this pattern and the greater rate of over-eruption with size may relate to the high degree of tooth wear experienced in this specialist osteophage or bone-eating species (2).

Over-eruption results in increasing diameter as well as height as the conical tooth emerges from the maxilla or mandible, with less increase in the medio-lateral axis than the other two dimensions. The reduced increase in medio-lateral diameter relates to canine shape and function in killing behaviour. Thylacines have sabre-like canines, quite narrow on the medio-lateral axis and broad in the anterio-posterior dimension, and thylacines were thought to kill prey small relative to their body size (2, 24, 25). Quoll canines are also sabre-like, but are more oval in cross-section, and are used in killing to penetrate and crush the back of the skull (2, 24). As a specialised osteophage, devil canines are rounded in cross-section (2), an adaptation that reduces the chance of canine fracture (22). Regardless of differences in killing function, for the canine teeth to retain strength with increasing height, it makes sense that they also increase in diameter.

What are the mechanisms of over-eruption and how do these influence the extent of over-eruption in different tooth types? In mammalian species with closed-rooted teeth, over-eruption or continuous eruption of teeth is reported to be a response both to tooth wear (11) and to the loss of the opposing tooth of an occlusal pair (12). I propose a third related mechanism in marsupial carnivores, that as an animal grows and body size increases the distance between the maxilla and mandible increases, functioning as an effective slow release of pressure from the occlusal paired tooth.

In the canine teeth of marsupial carnivores, the major mechanism driving over-eruption is possibly the gradual release of oblique occlusal pressure as the animal grows, with dynamic changes in tooth wear on the tip and sides of the tooth contributing to the changing occlusal pressure. The role of occlusion in the over-eruption could be complex. While the canine teeth do not have the complete occlusion seen in the molar teeth, the upper and lower canines do touch, sliding obliquely against each other during jaw movement. The strength of this mechanical interaction between upper and lower canine teeth results in substantial lateral wear facets in some individuals (Figure 1). This oblique contact between upper and lower canines may provide some occlusal pressure which gradually releases as the animal grows but with the increasing canine tooth diameter functioning to maintain it. The canine teeth in marsupial carnivores continue to over-erupt throughout life, suggesting primacy of a mechanism of low occlusal pressure although tooth wear plays an important role.

The significant contribution of tooth wear to canine over-eruption, in both marsupial and placental carnivore species, is suggested by the increasing amount of over-eruption with body size (which reflects age), found only in the two osteophagous species measured: the marsupial Tasmanian devil and the placental striped hyaena. Among carnivorous mammals, hyaneas and Tasmanian devils are the most specialised osteophages, with dental adaptations for consuming the hard parts of carcasses including bone (2, 22). Devils are ecomorphologically convergent with hyaenas, both evolving to occupy the bone-eating niche on different continents (2, 22). As a general pattern, the amount of canine over-eruption increases with tooth wear scores in wild devils, and also in quolls which are not bone-eaters, with some reduction at the highest wear score. In very old Tasmanian devils, the extent of canine tooth wear is such that the enamel cervical margin is near the tip of the canine tooth, with most of the crown having worn away. Canines in very old animals are prone to fracture (2), particularly if they also have wear facets on the sides, which reduce canine diameter and thus structural strength of the tooth.

The molar teeth in marsupial carnivores also over-erupt but to a much lesser extent than the canines and with substantial differences related to species’ body size and the eruption sequence. Molar over-eruption is most evident in the two larger species, the thylacine and the devil, and minimal in the two smaller species, the quolls. This may reflect differences in the size of the gap between the mandible and maxilla as individuals of these different species grow to mature size. In the two larger species, the extent of molar over-eruption decreases from anterior to posterior, correlating with eruption sequence (8) and the timing of eruption relative to the age and size of the individual. The greatest amount of over-eruption occurs in M1, with decreasing amounts in M2, then M3 and the least in M4. Carnivores need to have a functional set of teeth by the time they are weaned; canine teeth to kill and process prey and molar teeth with carnassial blade function to slice and consume meat. Unlike placental carnivores which have a specialised carnassial tooth (M1), each marsupial molar tooth (M1 to M4) has a carnassial blade and functions as the meat-slicing tooth as it sequentially erupts, with the functioning carnassial blade being in the same biomechanical position as the placental carnassial tooth 50% along the jaw (2, 4, 22), which is an optimal position for the carnassial function (26). The first and second molars (M1, M2) erupt prior to weaning when the denned young are eating solid food and are fully erupted by the time the juvenile marsupial carnivore is weaned, with M4 completing eruption within six months of independence. Sequential eruption means that M1 has fully erupted into the very small jaw of the dependent juvenile, while M4 erupts at a time when the animal is two thirds grown, the latter a similar body size when eruption of successional teeth takes place in placental carnivores. Thus, M1 will be exposed to a much greater increase in the distance between mandible and maxilla than M4 as the animal grows to mature size. In both the quoll species, for example, M1 increases with body size but M4 doesn’t.

The molar teeth in marsupial carnivores generally increase in height with body size as juvenile grow to maturity, but the life-long patterns across the molar tooth row vary among species, reflecting a combination of over-eruption and tooth wear, and thus species trophic ecology. The contribution of over-eruption to total tooth height is greater in the anterior molars which erupt first (M1, M2) than in the posterior molars which are last to erupt (M3, M4). In all marsupial carnivore species, molar height appears to reduce at the greater wear scores.

Because molar teeth have complete occlusion, over-eruption has a clear mechanistic role in compensating tooth height in response to growth in body size, to tooth wear (the amount of enamelled crown that has been worn away) and even to the loss of an occlusal partner (molar tooth fracture is rare in these carnivores 24). In adult wild great apes, for example, total tooth height does not vary with wear stage or age (11). The observed patterns of molar tooth height, over-eruption and wear across tooth number and species reflect tooth function, trophic ecology and ontogenetic differences between marsupial and placental carnivores.

Molar tooth wear is related to tooth function, being minimal in the carnassial teeth which need to stay sharp to slice meat effectively, and high in the bone-cracking teeth of specialised osteophages. The carnassial teeth are positioned well back in the jaw in both placental and marsupial carnivores (50% along the jaw length 4) and remain sharp during life in most species. Bone is processed further forward in the jaw, on P3 (premolar) in placental hyaenas (22) and M2 in marsupial devils (2), the two specialist osteophages in the study. This explains why M2 and also M3 in devils show extensive wear during life, actually decreasing in height. Even small devils eat bone and tooth wear, particularly on M2, is evident even in the first subadult year of independent life. By contrast, thylacines lack bone-eating specialisations and are thought to have killed prey small relative to their body size (2, 24, 25)). Thylacine museum specimens have low wear scores on all teeth and no individuals had a wear score on M4 greater than zero.

An unexplored potential consequence of tooth over-eruption is the abrasion strength of the exposed section of tooth beyond the enamelled proximal crown. Enamel structure is strongly correlated with tooth function and microwear on teeth is attributed to mechanical stresses of shearing and compression (27). In the larger marsupial carnivore species, devils and thylacines, but not in quolls, the premolar and molar teeth contain more enamel types than the incisors and canines, reflecting their function in shearing and grinding rather than grabbing and holding prey (28). In devils, compared with thylacines, there is less differentiation in enamel structure between the premolar and molar teeth, probably reflecting adaptation to their bone-eating diet (28). Placental osteophagous species, e.g. hyaenas, also have a more complex enamel structure (29). The question is whether the structural integrity of the exposed over-erupted tooth section that lacks enamel is compromised. This may not be the case if the microwear on the teeth is primarily on the leading edges of the canines and the molars (27), which remain protected by enamel.

This study has addressed an unresolved question in understanding the consequences of different ontogenetic patterns of tooth replacement in mammals, whether the jaw of species with a monophyodont tooth set are sufficiently large to accommodate adult-sized teeth at weaning when the animal needs functional dentition (3). Analysis of the ontogeny of changes in canine and molar tooth size, wear and eruption amongst the extant and recently extinct species of Australian dasyuromorph marsupial carnivores, that have monophyodont anterior dentition, reveals a mechanism of tooth over-eruption that circumvents the potential “constraint” of the monophyodont condition. Over-eruption, particularly of the conical-shaped canine teeth allows for a smaller canine to be fully functional in the jaw of a newly weaned juvenile and for the length and diameter of the tooth to increase as the animal grows to mature body size. The mechanisms that produce over-eruption in marsupial carnivore teeth appear similar those operating in other mammals, those being a consequence of tooth wear and loss of occlusal partner, with the variation that over-eruption may respond to the changing occlusal pressure as mandible and maxilla move further apart with growth in body size from juvenile to adult. Over-eruption in marsupial carnivores is substantially greater than that in ecologically equivalent placental carnivores which have diphyodont tooth replacement. Future research could explore this phenomenon across a broader range of marsupial and placental mammals exhibiting different patterns of tooth replacement.

## Acknowledgments

I thank Leon Barmuta for statistical assistance, Lars Werdelin, Richard Burn-Murdoch and Gordon Sanson for comments on an earlier draft; and the following museum curators and directors for assistance: Kathryn Medlock (Tasmanian Museum and Art Gallery), Tim Kingston (Queen Victoria Museum), Lina Frigo and Joan Dixon (Museum of Victoria), Linda Gibson and Sandy Ingleby (Australian Museum), Lynette Queale (South Australian Museum) and Tamar Dayan (Tel Aviv Museum). Data on wild animals were collected under Animal Ethics Permits numbers F.BTZ.11.00 (Australian National University) and A0009086 (University of Tasmania).

