## Supplemental Figures 1 and 2 for "Over-eruption in marsupial carnivore teeth: compensation for a constraint"

(A)

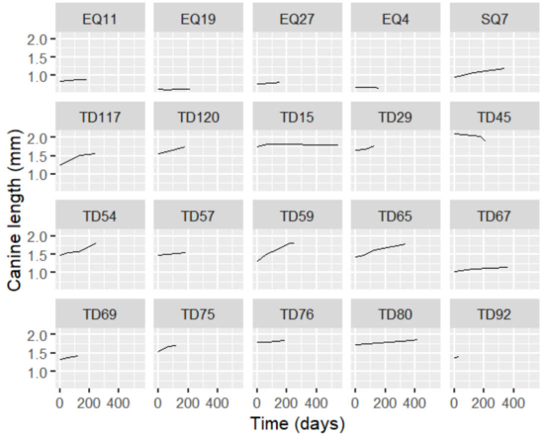


(B)

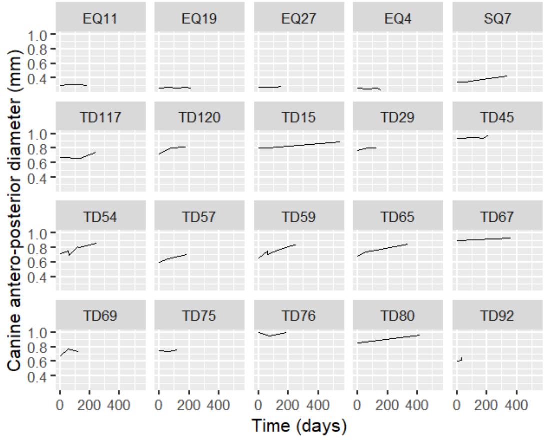


(C)

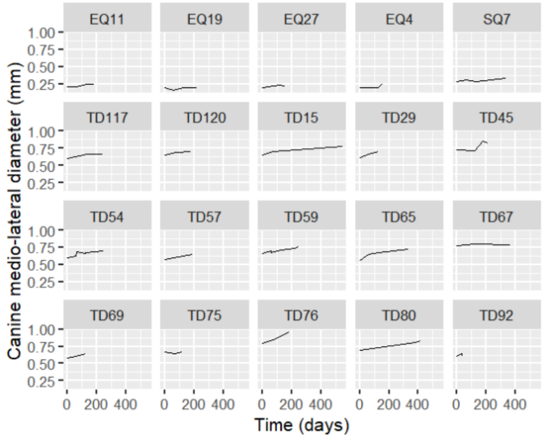


Supplementary Figure S1. Changes in canine dimensions with time in wild, living Australian dasyuromorph carnivores: total canine length (A); antero-posterior diameter (B); and medio-lateral diameter (C). cnl = right canine length (mm), measured from the apex of the gum line to the tip; apd = antero-posterior diameter and mld = medio-lateral diameter of the canine tooth measured at the gum line. Days = the number of days since the first capture of the individual at the study site at Cradle Mountain National Park, Tasmania. The individual figure titles include species code and individual identification number. EQ = eastern quoll, SQ = spotted-tailed quoll, TD = Tasmanian devil.

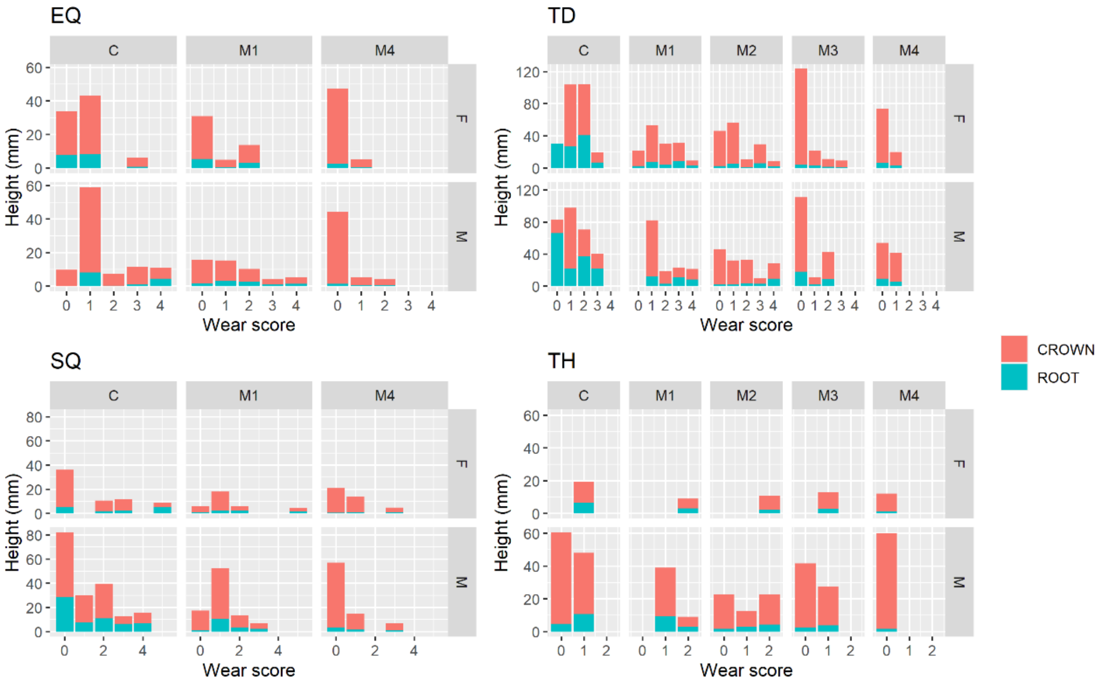


Supplementary Figure S2. Canine and molar tooth height for each tooth wear stage in marsupial thylacines, Tasmanian devils, spotted-tailed quolls and eastern quolls, wild-sourced from museum collections. The stacked bars separate the components of total tooth height that comprise the enamelled crown and the exposed root where the tooth has over-erupted. Tooth types are: canine (C), and molars 1 through 4 (M1, M2, M3, M4). Wear stages are 0 (no wear), 1 (tip or cusp worn), 2 (tip or cusp worn, dentine exposed), 3 (tip or cusp well worn, dentine exposed, less than half tooth height worn), 4(tip or cusp well worn, dentine exposed, more than half tooth height worn).
